# Path knockout: an analysis tool for connectome-driven circuit discovery

**DOI:** 10.64898/2026.10.31.763128

**Authors:** Ryan McGrath, Daisuke Hattori

**Affiliations:** Department of Physiology, UT Southwestern Medical Center, Dallas TX, USA; Department of Neuroscience, UT Southwestern Medical Center, Dallas TX, USA; O’Donnell Jr. Brain Institute, UT Southwestern Medical Center, Dallas TX, USA

## Abstract

Neural circuits in the brain transform sensory information into perceptual, emotional and motor output. The connectome – a synaptic-resolution wiring diagram of all neurons in a nervous system – provides a neuroanatomical blueprint for identifying different circuits that mediate specific computations. However, its sheer scale and complexity pose a significant challenge in extracting functionally relevant multi-synaptic pathways. Here we established *path knockout*, a connectome analysis method that identifies dominant multi-synaptic pathways. Path knockout uses the input fraction of each synaptic connection as a surrogate for the probability that information transfer succeeds across that synapse and calculates approximate probability, or *connection strength*, that at least one path succeeds among all multi-synaptic paths between defined source and sink neuron populations. It then quantifies the importance of each interneuron by assessing the impact of its removal on the connection strength. We demonstrate that path knockout outperforms other methods for pathway analysis and identifies known and novel interneurons in the *Drosophila* circuits. Using path knockout, we identified a convergent circuit motif linking sexually dimorphic aggression command neurons to a shared isomorphic visual pursuit pathway. Path knockout is implemented such that it runs in minutes on a standard computer. Thus, path knockout provides a scalable and generalizable framework for connectome-driven circuit discovery.

## INTRODUCTION

A fundamental goal of neuroscience is to understand how the brain transforms sensory inputs into perceptions and actions. This requires identifying neurons that participate in each behavior and understanding how information propagates through a network of neurons connected by synapses. Significant advances have been made in understanding how sensory systems detect and represent external stimuli and how motor outputs control muscles. However, identifying which pathways that comprise multi-synaptic strings of interneurons perform intermediate computations critical for the sensorimotor transformation remains a central challenge.

The connectome – a synaptic-resolution wiring diagram of all neurons in a nervous system – is an essential resource that provides a neuroanatomical foundation to identify different circuits underlying behaviors. It allows tracing potential information flow, generating hypotheses for circuit functions and rationally designing experiments and interpreting their results. The first connectome was determined for the *C. elegans* nervous system by reconstruction of serial-section electron micrographs (302 neurons, 7,000 synapses)^1^. This dataset has played a significant role in dissecting circuit mechanisms underlying worm behaviors. Later-determined worm connectomes have further provided foundational understanding of behavioral differences between sexes and between species^2,3^. The connectome comes with several limitations^4,5^: it is a snapshot of synaptic connectivity without dynamical information, does not contain the information of synaptic sign (but see ref^6^), and is also blind to the humoral or extrasynaptic effect of neuromodulation. Despite these limitations, the success enjoyed by the worm and fly neuroscience community demonstrates that the knowledge of synaptic connectivity offers a far-reaching advantage in understanding the architecture of neural circuits and the computation they perform.

Recent advances in volumetric electron microscopy, expansion microscopy and automated image analyses^7–9^ have led us into the era of connectomics. The growing list of available connectomes includes the entire *Drosophila* larval brain (3,016 neurons, 548,000 synapses)^10^, the entire adult female *Drosophila* brain (139,000 neurons, 50 million synapses)^11–13^ and the adult male *Drosophila* central nervous system (166,000 neurons, 46 million synapses)^14^. Extending connectomics to vertebrate, recent studies generated a connectome of a larval zebrafish brain (180,000 neurons, 30 million synapses)^15^ and a connectome for a volume of mouse visual cortex (200,000 neurons, 500 million synapses)^16^. Together, these datasets represent an extraordinary and rapidly expanding opportunity to systematically link neuroanatomy to function. Realizing the full potential of connectomes in identifying important pathways requires analysis methods capable of handling the sheer scale and complexity to extract biologically plausible circuit hypotheses.

A complete connectome is simply an exhaustive catalogue of synaptic connections across all neurons and thus the map of functional circuits needs to be extracted (**Fig. 1A**). The number of pathways that connect any pair of neurons grows exponentially with the number of interneuron layers and rapidly reaches a scale that is intractable for manual tracing. Two computational approaches have been used to discover multi-synaptic pathways connecting specific source (upstream) and sink (downstream) neurons; shortest-path analysis^10^ and maximum flow (maxflow) analysis^14^ (**Fig. 1B**). However, both approaches suffer from limitations in their ability to identify canonical neural pathways. Shortest-path analysis finds one pathway connecting source and sink with minimum number of synaptic “hops”, or interneuron layers, thereby prioritizing a shorter path even those with miniscule synaptic connections over a longer path with abundant connections. In addition, by returning a single “optimal” path, this algorithm does not allow us to compare alternative routes for follow-up experiments. Maxflow analysis quantifies the maximum signal that can be transmitted from a source to a sink across many polysynaptic pathways. However, it suffers from a biologically invalid assumption, rigid flow conservation, which requires the quantity entering a node (i.e., neuron) to be equal to the quantity exiting it. This effectively treats interneurons as passive cables that simply relay input signals, ignoring the nonlinear transformation each interneuron performs. Thus, there is a need for biologically grounded scalable computational approach to systematically identify and rank multi-synaptic pathways.

**Figure 1.**
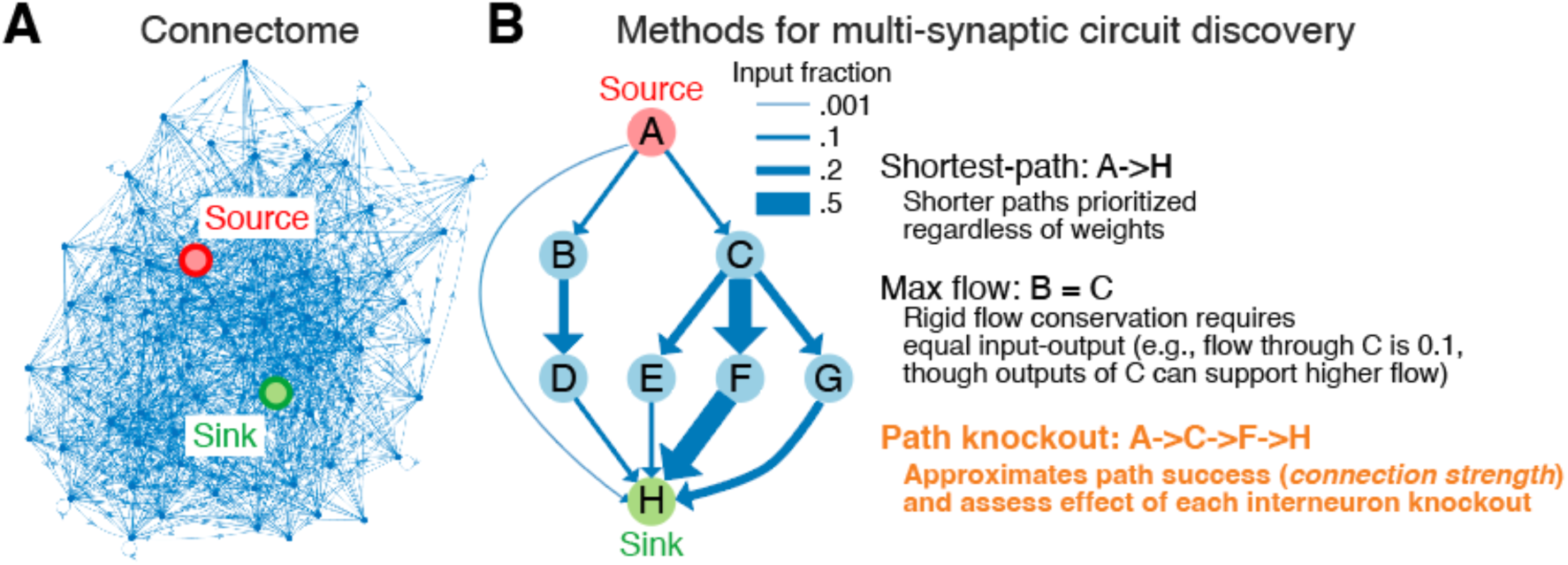
Path analysis concept. **(A)** Diagram of a connectome network after grouping by neuron type and reducing to nodes 2 hops upstream of sink and 2 hops downstream of source. **(B)** Conceptual diagram of network analysis given a target source and sink. Shortest path finds a key path regardless of edge weights. Maxflow finds key interneurons yet suffers from single edge bottlenecks. Path knockout evaluates total paths through each neuron with respect to edge weights along path.

In this study, we established a novel connectome analysis approach – termed *path knockout* – that bridges this gap in connectome-driven circuit discovery (**Fig. 1B**). Path knockout uses the input fraction of each synaptic connection as a surrogate for the probability that information transfer succeeds across that synapse and calculates approximate probability, or *connection strength*, that at least one path succeeds among all the paths that connect a source to a sink. This formulation of connection strength rewards circuits with many redundant parallel paths and treats each interneuron as an active contributor to – rather than a passive conduit of – information transmission. Path knockout then systematically assesses how removal of each interneuron reduces this overall connection strength, rank-ordering interneurons by their importance for source-to-sink information transfer, thereby revealing a canonical circuit linking source and sink. We employ the *Drosophila* model for development and validation of this method, as it provides an ideal system: in addition to multiple adult brain connectomes, decades of fly neuroscience have identified functional circuits mediating different behaviors, providing a robust set of ground truth against which path knockout results can be benchmarked. Furthermore, the unparalleled toolkit available for cellular-resolution genetic and neural circuit manipulation and recording^17–23^ enables experimental testing of circuit hypotheses generated by path knockout. We envision that the path knockout method will serve as a tool that bridges the gap between neuroanatomical wiring diagram and functional circuit hypotheses, facilitating connectome-driven circuit discovery.

## RESULTS

### Mathematical framework of path knockout

We represent a connectome as a directed weighted graph *G* = (*V*, *E*, *W*), where *V* is the set of neurons (or nodes), *E* the set of directed synaptic connections (or edges), and *W* the set of synaptic (or edge) weights. The weight *w*(*i*, *j*) of edge (*i*, *j*) is defined as the input fraction – the proportion of the total synaptic input onto postsynaptic neuron *j* that originates from presynaptic neuron *i*:

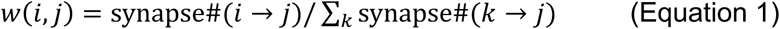

We interpret *w*(*i*, *j*) as a surrogate for the probability that a signal is successfully transmitted from neuron *i* to neuron *j* across that synapse (regardless of the sign of synapse; see Discussion). For a multi-synaptic path *π* = (*v*_0_, *v*_1_, … , *v_n_*) connecting source *S* = *v*_0_ to sink *Sk* = *v_n_* through *n* − 1 interneurons, the path success probability is the product of transmission probabilities across all edges:

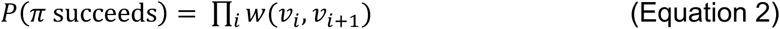

Let Π = (*S*, *Sk*) denote the set of all simple directed paths from *S* to *Sk*. Assuming path independence (see below), the *connection strength*, *CS*(*S*, *Sk*) – the probability that at least one path successfully transmits information from source to sink – is:

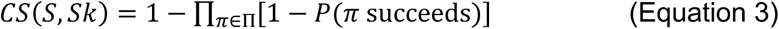

Path knockout of interneuron *n* is performed by removing *n* and all its edges from *G* to yield the subgraph *G*^−*n*^ and recalculating connection strength on this reduced graph. The *knockout (KO) index* of interneuron *n* is then defined as:

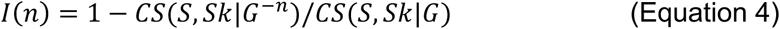

The KO index is a normalized ratio bound by 0 and 1: *I*(*n*) = 0 when removing *n* has no effect on connection strength (dispensable neuron), and *I*(*n*) = 1 when removing *n* reduces connection strength to zero (essential neuron). Interneurons are rank-ordered by *I*(*n*), with higher scores indicating greater importance to source-to-sink information transmission.

### Practical implementation of path knockout

We implemented path knockout (**Fig. 2A**) first using the *Drosophila* hemibrain dataset^7^. We employed three strategies to make path knockout computationally tractable without sacrificing biological relevance. First, we extract a subconnectome comprising all neurons within 2 hops downstream of the source neuron population and all neurons within 2 hops upstream of the sink neuron population. This subconnectome includes all true 5-hop connections, more than the average shortest-path length connecting any pairs of neurons (4.42) in the full adult female brain connectome (FAFB)^24^. Second, we group neurons by cell type into a single node and define the edge weights as the sum of all synaptic connections from cell type *i* to cell type *j*, divided by the total synaptic input onto cell type *j* from all sources, thereby preserving the input fraction interpretation of *w*(*i*, *j*) at the cell-type level. Third, paths within a subconnectome are identified using the MATLAB `allpaths` function. The function is used to identify only simple paths up to 5 hops with no recurrence, focusing on extracting the feedforward pathways from source to sink. Through these optimizations of analysis parameters, path knockout achieves full analysis of each subconnectome comprising hundreds to thousands of neuron types within a few to ten minutes with a standard desktop computer with 16GB RAM. The analysis pipeline then returns the overall connection strength of all paths connecting the source and the sink, a trimmed circuit graph containing top-ranked interneurons by KO index, and KO index for each interneuron in the subconnectome (see e.g., **Fig. 2B**).

**Figure 2.**
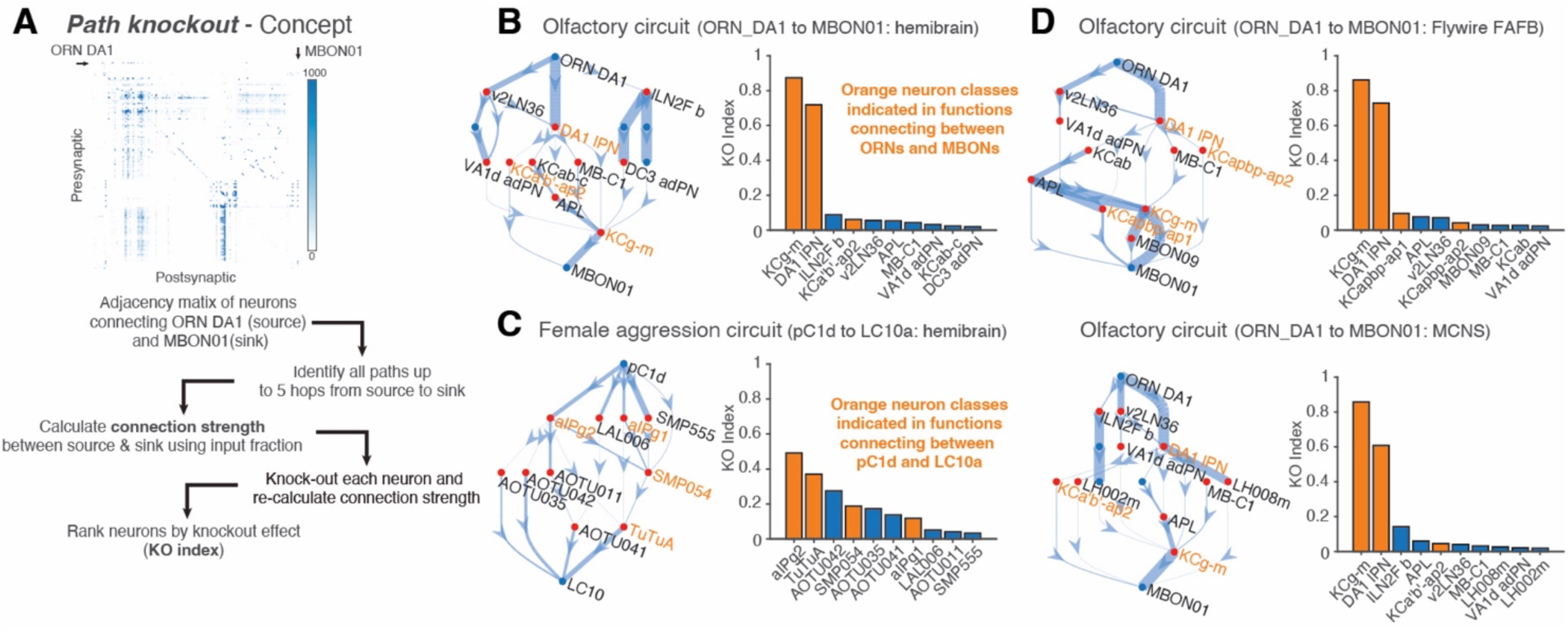
Path knockout finds key interneurons in multiple networks across connectome datasets. **(A)** Path knockout concept. Connection strength is determined for original network and all sub networks with one node knocked out. Nodes are sorted based on total reduction in connection strength post knockout. **(B)** Results of path knockout performed on olfactory circuit, ORN DA1 to MBON01 using the hemibrain dataset. Left; trimmed down graph showing the top 15 paths from source to sink. Top 10 nodes found with path knockout are highlighted in red. Orange text denotes known canonical interneuron groups. Edge width indicates input fraction. Right; bar graph of top 10 nodes and their respective knockout (KO) index. Orange highlights known canonical interneuron groups. **(C)** Results of path knockout for female aggression circuit (pC1d to LC10a) using the hemibrain dataset. **(D)** Results of path knockout for olfactory circuit using the FAFB (top) and the MCNS (bottom) datasets.

### Path knockout recovers known interneurons in olfactory circuit and female aggression circuit

As a proof of principle, we first applied path knockout to the olfactory circuit, one of the best-characterized circuits in the fly^25^ (**Fig. 2A**). Each odorant receptor neuron (ORN) expresses a single odorant receptor and all ORNs expressing the same receptor converge on a specific glomerulus in the antennal lobe, where they synapse with projection neurons (PNs) and local interneurons (LNs). Uniglomerular PNs restrict their dendrites to a single glomerulus, whereas multiglomerular PNs arborize across multiple glomeruli. While LN dendrites and axons are confined within the antenal lobe, PN axons transmit the olfactory information to two higher brain centers, the mushroom body and the lateral horn. In the mushroom body, PNs synapse onto the principal mushroom body neurons, Kenyon cells (KCs), which in turn synapse with 34 classes of mushroom body output neurons (MBONs). There are at least three distinct KC classes (gamma, alpha’/beta’, and alpha/beta) and each MBON type receives presynaptic input from one or two of these KC classes. Source neurons were defined as a class of ORNs expressing the Or67d receptor (a receptor for a male pheromone, cVA)^26^ that innervate the DA1 glomerulus (ORN_DA1) and sink neurons were defined as a class of MBONs (MBON01)^27^ that receives input from the gamma and alpha’/beta’ KCs. Path knockout identified the uniglomerular PNs of the correct glomerulus (DA1_lPN) and the gamma and alpha’/beta’ KCs as the top-ranked interneurons by KO index, consistent with their role as the obligate relay (**Fig. 2B**). Thus, path knockout identifies the dominant interneurons in the 3-hop pathway connecting the ORNs to the MBONs.

We next applied path knockout to a central brain circuit – the female aggression circuit. Female aggression in *Drosophila* is manifested as a sequence of pursuit and contact behaviors (e.g., headbutting and fencing) gated by the external environment, such as the presence of food and egg-laying sites, and by the internal state, such as hunger and mating status. Previous studies identified two central brain neuron classes, pC1d and its downstream target aIPg, that drive aggressive behavior in females^28^. pC1d neurons integrate internal states relevant for aggression, whereas aIPg neurons gate the LC10a visual processing pathway that promotes pursuit of a moving fly^29–33^ through 3-hop disinhibitory pathway containing two inhibitory interneurons, SMP054 and TuTuA. Source neurons were defined as pC1d neurons and sink neurons were defined as LC10a visual projection neurons. Path knockout of the pC1d-LC10a subconnectome identified all the interneurons with known function in this 4-hop pathway (aIPg, TuTuA and SMP054)^28,32^ as well as novel interneurons with no prior functional characterization (**Fig. 2C**).

Given these initial successes, we expanded path knockout to other *Drosophila* adult connectome datasets; FAFB (full adult female brain)^11–13^ and MCNS (male central nervous system)^14^. Path knockout applied to the ORN_DA1-to-MBON01 pathways in the FAFB or MCNS dataset revealed KO index rankings aligning with those obtained in the hemibrain dataset with equivalent computation time (**Fig. 2D**). These results demonstrate that path knockout is capable of identifying and rank-ordering canonical interneurons within multi-synaptic pathways.

### Validation of the path independence assumption and computational parameters

We next examined the impact of the path independence assumption employed for path knockout. The formulation of connection strength (**Equation 3**) makes a convenient assumption that each path from source to sink fails independently, but this is biologically incorrect as paths sharing one or more edges are not truly independent. We performed Monte Carlo simulations in which signals are stochastically propagated through the subconnectome to obtain empirical connection strength estimates that do not require the independence assumption (**Fig. 3A**). We then compared the connection strengths for all paths that traverse a given node that are obtained by 5-million Monte-Carlo trials with those that are analytically obtained by path knockout. The connection strengths were highly concordant between the two methods in each of the four subconnectomes analyzed (**Fig. 3B**). To evaluate the rank orders more quantitatively, we performed rank-order rescue by attempting to restore rank-order by shifting Monte Carlo probability within bootstrap-determined 95% confidence interval (**Methods**). This analysis found that all top-ranked nodes were rescued in each of the four subconnectomes analyzed (**Fig. 3B**). These results demonstrate that, although path independence assumption introduces approximation error in the absolute connection strength value, it does not affect the KO index rankings obtained by analytic path knockout equations.

**Figure 3.**
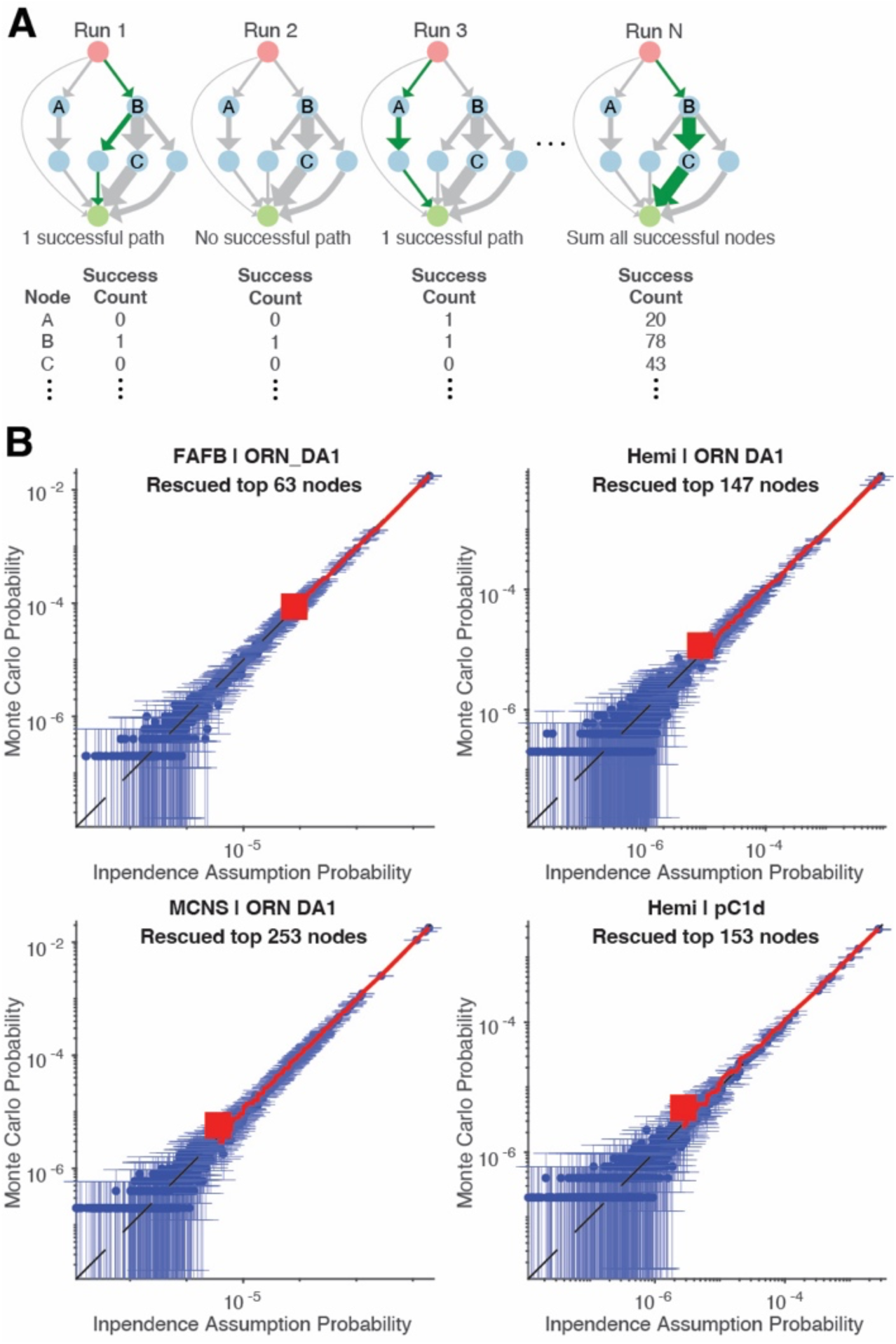
Validation of path independence assumption for path knockout. **(A)** Monte Carlo simulation concept. Per iteration (run), nodes on successful paths are recorded and a running total for each node is kept across runs. Total node successes and number of runs are then used to estimate the probability that at least one path was successful that contains a node without assuming path independence. Paths that account for up to 99.5% of overall connection strength in each subconnectome were used for Monte Carlo analysis for computational efficiency (**Methods**). **(B)** Comparisons of the probability that at least one path was successful that contains a node with (used for path knockout) or without (Monte Carlo simulation) independence assumption in the four subconnectomes. Blue dots, nodes (interneurons); error bars, 95% confidence interval of Monte Carlo probability; redlines, the top N nodes that are rank-rescued.

### Comparison of path knockout performance with other analyses

We next compared the performance of path knockout with other methods. Weighted shortest-path analysis returns a with minimal hop lengths with minimal distance across nodes connecting between a source and a sink. When used for connectome with edge distance defined as complement of input fraction (i.e., 1 − *w*(*i*, *j*)), weighted shortest-path analysis can identify the strongest pathway. The primary limitation of this shortest-path analysis is that it prioritizes shorter paths with miniscule edge weight over longer paths with more abundant weight. Combining this algorithm with weight thresholding, however, may potentially rectify this limitation. We devised a method to iteratively identify shortest-paths linking a source to a sink with increasing edge thresholds (**Fig. 4A**; **Methods**). This iterative shortest-path search applied to the olfactory circuit linking ORN_DA1 to MBON01 correctly identified the two dominant interneuron classes (DA1_lPN and KCg-m) as well as the dominant pathway through these neurons (**Fig. 4B**). However, it entirely missed another important interneuron class, KCa’b’, which was found in path knockout (**Fig. 2D**), and it also found neurons likely unrelated to the feedforward pathways (e.g., PAM01). Applying this method to the female aggression circuit connecting pC1d with LC10 also found dominant interneuron classes and the correct 4-hop pathway (**Fig. 4C**). However, each canonical interneuron class was found in different shortest paths sporadically without apparent rank orders, and the correct pathway was buried as the 6^th^ best shortest path. These results indicate that, while this iterative shortest-path analysis can identify dominant interneurons and pathways, paths found are greatly affected by threshold chosen. Moreover, the lack of quantitative information, other than the threshold, that describes each path limits its utility in comparing alternative pathways.

**Figure 4.**
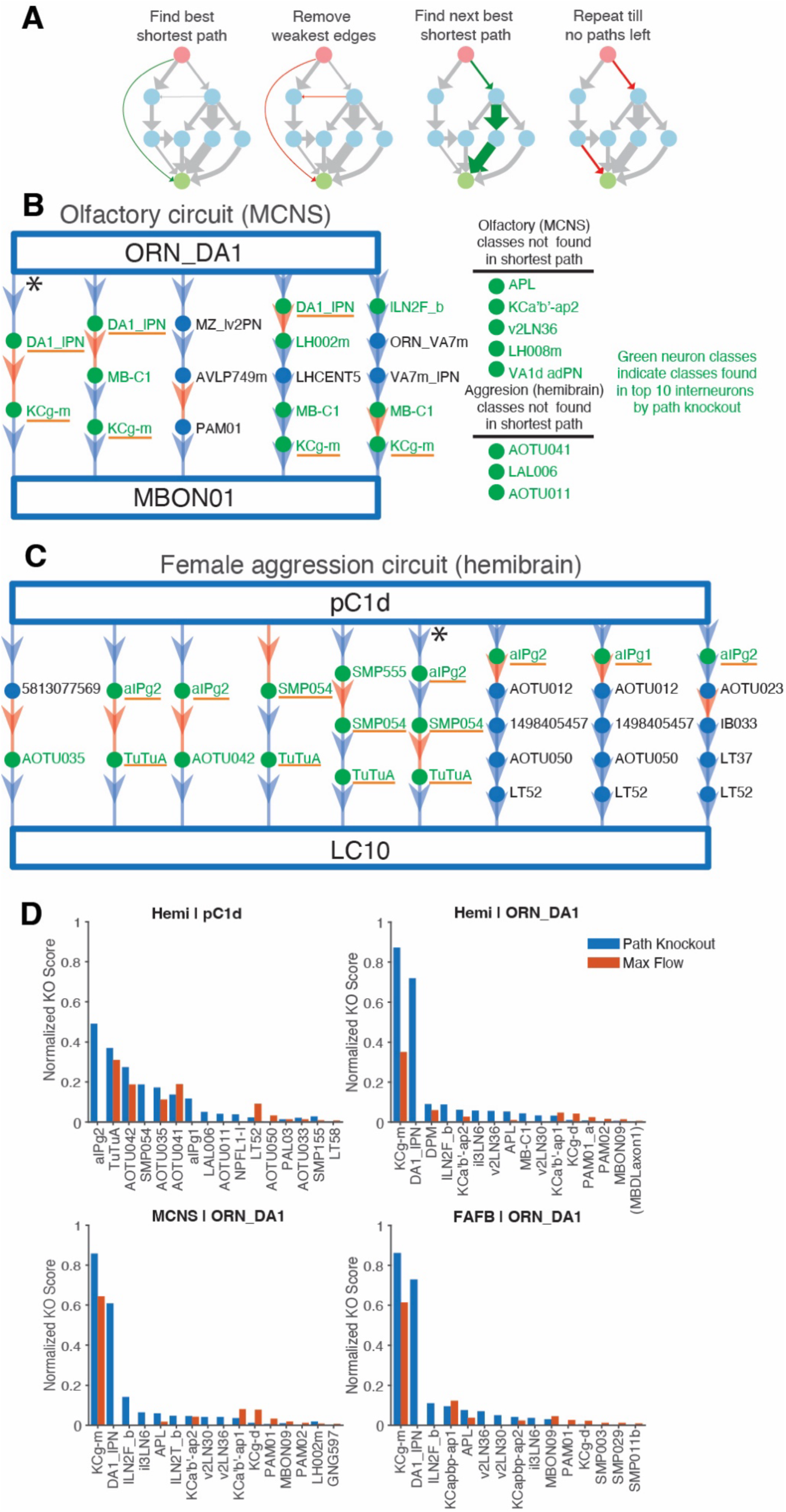
Comparison of path knockout to shortest path and maxflow analysis. **(A)** Shortest-path analysis with iterative edge thresholding. A single shortest path (weighted) in each graph was identified, edges with weight equal to or smaller than the smallest edge in this path were removed, and the shortest-path algorithm was applied to the new thresholded graph. **(B)** Results of iterative shortest-path analysis applied to ORN_DA1-to-MBON01 circuit. The paths are ordered from left to right by increasing edge thresholds (i.e., the left most path is the shortest path of the intact subconnectome). Edges with the smallest weight in each path (i.e., to be removed in the next iteration) are indicated in red; top 10 path knockout interneurons by KO index are in green with those not found in the shortest-path analysis in the list on the right; interneurons functionally validated by prior studies are underlined in orange; and the functionally validated canonical pathway is indicated by asterisk. Only paths with a hop number fewer then 7 are shown. **(C)** Results of iterative shortest-path analysis applied to pC1d-to-LC10 circuit. **(D)** Comparison of normalized maxflow knockout index with path analysis knockout index across four circuits.

Maxflow analysis quantifies the signals transmitted from a source to a sink through all paths and thus is advantageous over shortest-path method. To test the ability of maxflow to identify canonical paths, we implemented maxflow with iterative interneuron removal (*maxflow knockout*), just as path knockout (**Methods**). The reduction in the maxflow per node removal was then quantified and normalized to match the formulation of path knockout KO index. This effectively allows us to compare the KO rankings obtained by maxflow vs path knockout, providing a way to assess the overall accuracy of using maxflow value vs connection strength in inferring the quantity of signals that can be transmitted through a subconnectome. We applied this analysis to the four subconnectomes used for our proof-of-principle analyses. Comparison of the KO rankings revealed that maxflow knockout missed some of the major interneurons in these circuits (e.g., DA1_lPN in the olfactory circuit and aIPg and SMP054 in the aggression circuit; **Fig. 4C**). This is likely because maxflow is an algorithm that finds a rate-limiting bottleneck in a graph, which is insensitive to interneurons that contributes to connection strength through many parallel paths. Thus, these analyses demonstrate the utility of connection strength in quantifying overall information transmission through multi-synaptic paths and the performance of path knockout in identifying dominant pathways linking a source and a sink.

### Source screen identifies dominant olfactory inputs and their interneuron pathways to pC1

Connection strength quantifies the aggregate influence of one neuron population on another across all multi-synaptic pathways. We exploited this property to identify strong source-sink combinations linking different source neurons in a layer (e.g., sensory neuron types) with a downstream neuron. We performed this source screen in the olfactory system by computing connection strength in the hemibrain dataset from each of 48 ORN classes to pC1 neurons – a central command node for female social behavior. This analysis identified that ORN_DA1 provides the largest connection strength to pC1 among all ORN classes (**Fig. 5A**). Examination of the KO index indicates a multiglomerular projection neuron type (M_lvPNm45) as the largest contributor to the connection strength of this pathway (**Fig. 5B**). Notably, previous functional studies found that optogenetic activation of these lvPNs induce female receptivity, serving as a crucial relay connecting cVA detection to social behavior^34^, validating the functional relevance of our analysis results. Our analysis also identified ORN_V and ORN_DP1m as additional olfactory channels with strong structural influence on pC1 neurons. The V glomerulus responds to CO2, a component of the *Drosophila* stress odor^26^, raising the possibility that stress odorant modulates female social behavior through a multi-synaptic path to pC1. Thus, source screen can effectively identify specific source neuron populations with potential functional impact on a downstream neuron, helping generate experimentally testable circuit hypothesis.

**Figure 5.**
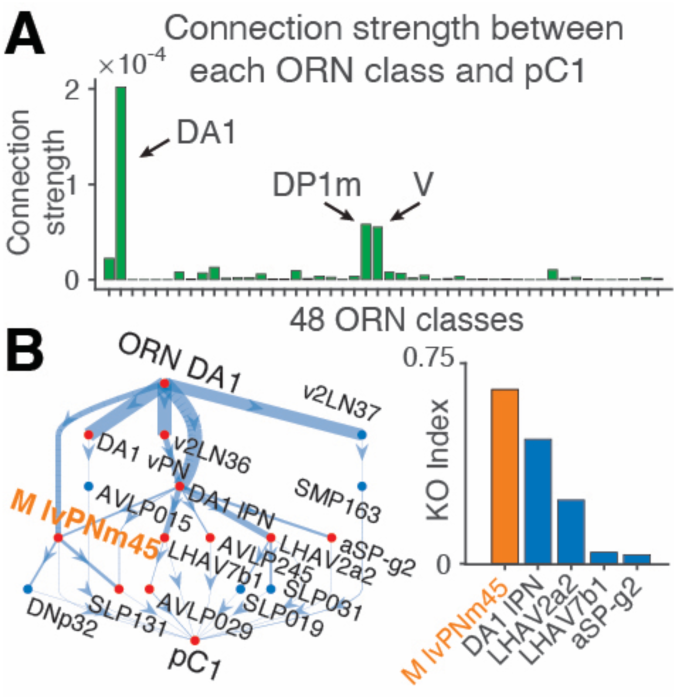
Source screen identifies dominant olfactory input channel and its pathway to the hub for female social behavior. **(A)** Connection strength from each ORN type to pC1 neurons. Hemibrain dataset. **(B)** Results of path knockout for ORN_DA1-pC1 pair that exhibited the largest connection strength. Orange highlights an atypical PN types found in a previous functional study (ref34).

### Path knockout identifies a sexually isomorphic pathway for modulating visually guided behavior during aggression

Finally, we applied path knockout to examine pathway structures between two layers of neurons, focusing on central brain circuits mediating social behaviors. pC1 neurons are sexually dimorphic central neurons that integrate multisensory conspecific signals and drive diverse social behaviors^35,36^. In females, five pC1 classes (pC1a-pC1e) regulate aggression and mating receptivity^28,37–40^, while in males, an expanded population of 23 pC1 supertypes similarly regulate distinct social behaviors like aggression and courtship^41–45^. Activation of different pC1 neurons likely induces distinct social arousal state that then shapes the activity of downstream effector circuits controlling specific behavioral features. One such behavioral feature is visually guided locomotion: for example, flies engaging in courtship or aggression pursue or flee from another fly. Visually guided behaviors depend on some 40 classes of visual projection neurons (VPNs) that convey information about distinct visual features computed in the optic lobe into the central brain^29^, and pC1 neurons may shape VPN activity to gate visually guided locomotion during social arousal. Indeed, activation of a courtship-promoting pC1 subclass (known as P1a) was shown to regulate visual pursuit of a conspecific target through modulation of response gain in a VPN class, LC10a^31^. Whether pC1 subclasses that drive different social behaviors recruit VPN circuits equally, or whether VPN connectivity reflects a subclass-specific behavioral function, remains unknown.

To address this, we computed connection strength from each of 28 pC1 subclasses across both sexes to each of 45 VPN classes using path knockout as a layer screen (5 hops). This generated a pC1-by-VPN connection strength matrix that captured the polysynaptic structural influence of each pC1 subclass across the VPN channels. We also included TK^FruM^ neurons, a class of male-specific neurons co-expressing the neuropeptide tachykinin and the male-specific isoform of the transcription factor *fruitless* (FruM), whose activation induces a robust aggression^46^, because they represent a major male neuron type that drives aggression not belonging to the pC1 class. Clustering of this matrix revealed that pC1 subclasses segregate into two discrete groups (**Fig. 6A**). One cluster contains pC1d and pC1e (female) together with pC1_9, pC1_10, and TK^FruM^ (male), whereas the second cluster contains the remaining pC1 subclasses from both sexes. Notably, pC1d and pC1e are known to drive aggression in females^28,40,47^, and a recent study suggested that pC1_9 and pC1_10 may play analogous roles in male aggression based on their connectivity^44,32^. These results suggest that neurons involved in aggressive arousal may share a distinct polysynaptic connectivity pattern to VPNs, and that aggressive arousal circuits in females and males may recruit a similar set of VPN pathways to shape visually guided behavior.

**Figure 6.**
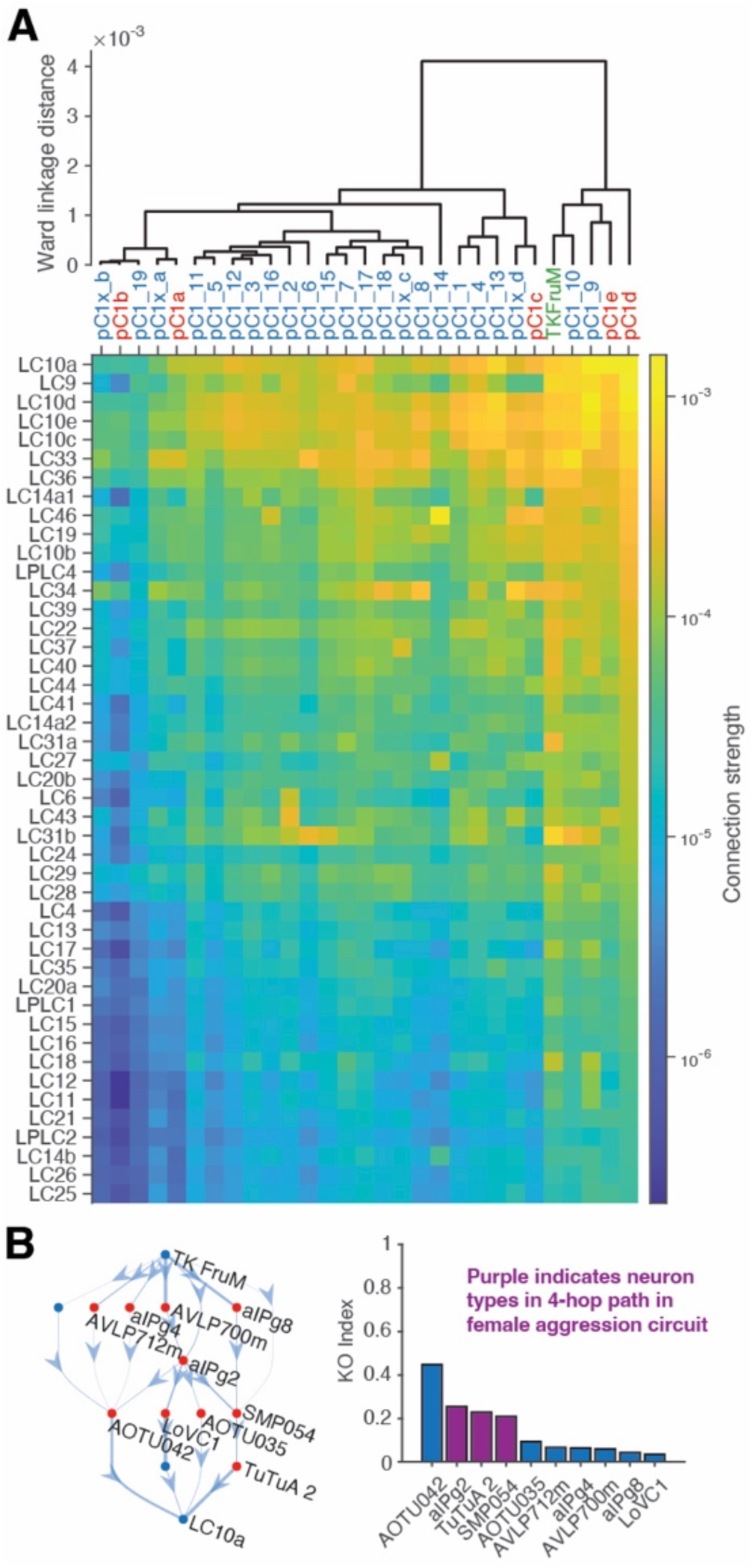
A layer screen between pC1 social arousal neurons and visual projection neurons identifies sexually isomorphic pathways linking aggressive arousal to visual pursuit. **(A)** Top, dendrogram showing clustering (Ward linkage) of connection strengths between each social arousal neuron class (pC1s: females, red; males blue; and TK^FruM^: green) and each visual projection neuron class (VPNs). Bottom, matrix of connection strengths. **(B)** Path knockout results for TK^FruM^-to-LC10a circuit (MCNS dataset).

Examining the specific pathways, we found that connection strengths from aggression-promoting neurons were highest to VPNs tuned to small moving objects, prominently including LC10a. Moreover, comparing interneurons with high KO index in the TK^FruM^-to-LC10a pathway in males with those in the pC1d-to-LC10a pathway in females revealed the same 4-hop circuit traversing aIPg, SMP054 and TuTuA neurons (**Fig. 6B** and **Fig. 2C**). This shared interneuron architecture indicates the existence of a sexually isomorphic circuit linking aggressive arousal to visual pursuit. Notably, this exact aIPg-SMP054-TuTuA-LC10a circuit in females has been shown to function in regulating overall aggression, raising the possibility that the male counterpart – connecting TK^FruM^ to LC10a through the same interneurons – plays a parallel role in male aggression.

Previous studies revealed a circuit motif in which upstream sexually isomorphic neurons (CAP) drive aggressive pursuit and downstream sex-specific neurons (fpC1 in female and MAP in male) dictate sex-specific attack behaviors^39^, illuminating a circuit logic in which an isomorphic central aggressive state affects dimorphic consummatory actions. Our results indicate that, in addition to isomorphic-to-dimorphic circuit motif, the opposite also exists, in which activation of sexually dimorphic central neurons (pC1d/pC1e in females, TK^FruM^ in males) converges onto an isomorphic downstream circuit mediating a common behavioral feature – visual pursuit – manifested during aggression in both sexes. This motif may enable transformation of aggression-promoting sensory input that may be dimorphic – like the presence of courtship target for males vs egg-laying patch for females – into isomorphic behavioral programs, illuminating a circuit logic that controls aggression. It will be interesting to examine whether connections from TK^FruM^ or pC1d to other VPN classes gate additional visually guided behavioral features, such as collision avoidance or looming-evoked escape.

## DISCUSSION

In this study, we established path knockout, a computational method that quantifies the contribution of interneurons to signal transmission between a source and a sink neuron population through multi-synaptic pathways in a connectome. By using connection strength as a surrogate for the probability that at least one path successfully transmits signal from source to sink and by assessing the effect of each interneuron removal to this probability, path knockout rank-orders interneurons by the importance of each neuron in the source-to-sink signal propagation, thereby helping identify dominant pathways without prior functional knowledge of the circuit. Applied to the well-characterized olfactory and female aggression circuits, path knockout correctly identified the known relay interneurons, validating the approach against ground truth. We then applied path knockout to previously uncharacterized circuits and found a convergent, sexually isomorphic circuit motif linking sexually dimorphic aggression command neurons (pC1d in female vs TK^FruM^ in male) to the LC10a visual pursuit pathway through a set of shared interneurons. Together, these results establish path knockout as a tool for discovering dominant multi-synaptic pathways from connectome data.

Several limitations exist for path knockout. First, by restricting the analysis to simple paths, path knockout excludes recurrent loops. Large-weight backward connections can significantly affect circuit dynamics: for example, feedback inhibition by a large inhibitory interneuron APL in the mushroom body is known to sparsen KC responses^48^, likely affecting the ORN-to-MBON signal transmission. The current path knockout framework does not capture the feedback-dependent contributions of an interneuron to source-sink signal transmission. Second, path knockout does not consider all paths within a subconnectome, as the path cap is required for computational tractability. Thus, connection strength always underestimates the true probability of transmission success. Third, the 5-hop ceiling excludes contributions of interneurons in longer hop pathways. Fourth, input fraction used here as a surrogate for signal transmission probability does not reflect the physiological efficacy of synaptic transmission. Moreover, path knockout is agnostic to the sign of synaptic connections and treats excitatory and inhibitory interneurons equivalently as contributors to connection strength. Finally, path knockout treats all synapses onto a neuron as functionally equivalent regardless of subcellular location, whereas synapses in different subcellular compartment, like dendrites or axons, distinctly contribute to the output of each neuron.

Despite these limitations, path knockout provides considerable utility as a tool for connectome-guided circuit discovery. It allows efficient identification of candidate interneurons across subconnectomes comprising hundreds to thousands of neuron types. This efficacy also enables layer-to-layer screen to find strong channels of connections between neurons in different processing layers. Finally, its light computational weight is well suited to serve as a companion tool for computationally intensive network simulation brain models that consider synaptic signs^49,50^, for example, by providing a ranked list of interneurons to be assessed for their significance in supporting simulated network activity. Thus, we envision that path knockout will provide a useful tool that bridges neuroanatomical connectivity analysis, computational simulation of network dynamics and experimental dissection of circuit function, facilitating connectome-driven neuroscience.

## METHODS

### Key Resources Table

| REAGENT or RESOURCE | SOURCE | IDENTIFIER |
|---|---|---|
| Software and algorithms |  |  |
| MATLAB (2023b, 2025b) | MathWorks | https://www.mathworks.com/ |
| Path Knockout Analysis Package | This paper | Github |
| neuPrint API | Scheffer et al. <sup>7</sup> | https://github.com/connectome-neuprint/neuprint |
| Other |  |  |
| neuPrint MCNS | Berg et al.<sup>14</sup> | https://male-cns.janelia.org |
| neuPrint hemibrain | Scheffer et al. <sup>7</sup> | https://neuprint.janelia.org |
| Flywire FAFB | Dorkenwald et al. <sup>12</sup> | https://codex.flywire.ai/api/download |

### Connectome Datasets

FAFB and MCNS connectome datasets were downloaded from respective online databases. The downloadable hemibrain dataset lacks some neurons such as ORN DA1 from the downloadable set. Therefore, we used the provided API for neuPrint server to download additional information from the live updated online hemibrain dataset. To avoid excessive calls to the API, we focused our attention on the neurons relevant to our current path scans by only retrieving connectivity of neurons downstream of targeted source and upstream of sink. We added the found neurons and their connections to our local dataset which we refer to as the extended hemibrain dataset. All analysis is performed on these local datasets. Local connectomes contain a list of body IDs, their group identification (neuron type), and their connectivity to other neurons in synapse count. As input into our path knockout analysis, we first trim connectivity to only include connections above a minimum synapse weight (default; greater than or equal to 5). Next, we group neurons on their group identification combining their connectivity. Input fraction is calculated on the synapse counts seen in this trimmed and combined dataset as the number of synapses pre- to postsynaptic neuron group divided by the total input into the postsynaptic neuron group.

### Path Knockout

To determine connection strength between a source and sink neuron types within a defined network, we first find all neurons two hops upstream from the sink and two hops downstream from the source, capturing up to all true 5-hop connectivity. We set the edges within this sub-network adjacency matrix (subconnectome) to be the number of synapses from neuron A to neuron B divided by the total synapses going into neuron B (input fraction). We assume the probability that an edge succeeds is equal to input fraction. We use MATLAB `allpaths` function to identify all simple paths up to 5 hops from source to sink within the subconnectome. We assume path independence and operationally define that the connection strength from a source to a sink is the probability of at least one path succeeds. We determine importance of individual nodes in the subconnectome by removing each node and recalculating connection strength on the new subconnectome. The reduced connection strength is normalized by dividing by the original connection strength of the network and then taking one minus the value gives knockout (KO) index.

### Monte Carlo

To evaluate the impact of the path independence assumption, we used Monte Carlo simulation of edge successes to estimate the real probability that at least one path from source to sink succeeds. For each run in the simulation, we sample the edges in the adjacency matrix to determine which edges succeed. If all edges along at least one path succeeds, that run is considered a success and the running total of successful nodes in the path is kept. Due to the small chance that any one path succeeds, Monte Carlo simulation is not feasible across all the paths found. We therefore limit the paths used for simulation to the paths that account for 99.5% of overall connection strength between a source and a sink, keeping the stronger paths. We also employ Poisson distribution instead of binomial for edges with weight less than 0.05 for computation efficiency, as Poisson produces equivalent results to bionomial for small probability and is more efficient. With these limitation, Flywire FAFB olfactory circuit used 195,572 (0.0095% of total) paths. neuPrint hemibrain olfactory circuit used 411,080 (0.02% of total) paths. neuPrint MCNS olfactory circuit used 97,693 (0.0043% of total) paths. neuPrint hemibrain aggression circuit used 261,939 (0.0129% of total) paths. We ran a total of five million simulations for each evaluated network. The impact of independence assumption is evaluated by attempting to rank-rescue node importance within the subconnectome. Connection strengths from source, sink, and intermediate node can be shifted within the 95% confidence interval of the Monte Carlo simulation estimated value.

### Shortest Path

To evaluate subconnectome using shortest paths, we converted a determined network into a MATLAB `digraph`. We took the complement of input fractions to be the edge weight (i.e., distance). We used MATLAB’s `shortestpath` function to find the current weighted shortest path from source to sink with respect to this distance, selecting the shortest path with smallest distance. Next, we find the edge with the greatest distance (smallest input fraction) in the identified shortest path and remove all edges with distance equal to or greater than the determined edge in the suconnectome. With the new `digraph`, we rerun `shortestpath` and repeat the removal of the edges till no paths between source and sink remain or the path length reaches 7 hops. We keep all identified paths for plotting.

### Max Flow

Max flow computes the greatest total flow that can be transmitted from a source to a sink with two constraints: the flow along each edge cannot exceed that edge’s capacity and the flow entering each node must equal the flow leaving the node. Input fraction was used as the edge weights and MATLAB’s `maxflow` function was used to determine the max flow from source to sink. To determine the importance of each node in supporting max flow, each node was individually removed from the network and max flow was recalculated. The reduced max flow values were normalized by dividing by the original max flow of the intact network and one minus the normalized value was used as the max flow knockout score.

## CODE AND DATA AVAILABILITY

Path knockout codes were written in MATLAB 2025b. A GitHub repository of all source codes for path knockout and data used in this manuscript with be made available once published on a public repository (access to current private repository available on request).

## ACKNOWLEDGEMENTS

We would like to thank Kenta Asahina, Wenhao Zhang and members of the Hattori lab for discussions. Avirut Mehta for support setting up initial code base. This research was supported in part by the computational resources provided by the BioHPC supercomputing facility located in the Lyda Hill Department of Bioinformatics, UT Southwestern Medical Center. This work is supported by NIH R01DK132705, UT Southwestern Endowed Scholars Program, and UT Southwestern O’Donnell Brain Institute.

## AUTHOR CONTRIBUTIONS

R.M. and D.H. conceived the project; R.M. wrote the source codes; R.M. and D.H. performed analyses and wrote the manuscript.

## DECLARATION OF INTERESTS

None to declare.

## Notes

### Competing Interest Statement

The authors have declared no competing interest.

## REFERENCES

1 White, J. G., Southgate, E., Thomson, J. N. & Brenner, S. The structure of the nervous system of the nematode Caenorhabditis elegans. Philos Trans R Soc Lond B Biol Sci 314, 1–340 (1986). 10.1098/rstb.1986.0056

2 Cook, S. J. et al. Whole-animal connectomes of both Caenorhabditis elegans sexes. Nature 571, 63–71 (2019). 10.1038/s41586-019-1352-7

3 Cook, S. J. et al. Comparative connectomics of two distantly related nematode species reveals patterns of nervous system evolution. Science 389, eadx2143 (2025). 10.1126/science.adx2143

4 Bargmann, C. I. & Marder, E. From the connectome to brain function. Nat Methods 10, 483–490 (2013). 10.1038/nmeth.2451

5 Scheffer, L. K. & Meinertzhagen, I. A. A connectome is not enough - what is still needed to understand the brain of Drosophila? J Exp Biol 224 (2021). 10.1242/jeb.242740

6 Eckstein, N. et al. Neurotransmitter classification from electron microscopy images at synaptic sites in Drosophila melanogaster. Cell 187, 2574–2594 e2523 (2024). 10.1016/j.cell.2024.03.016

7 Scheffer, L. K. et al. A connectome and analysis of the adult Drosophila central brain. Elife 9 (2020). 10.7554/eLife.57443

8 Chen, F., Tillberg, P. W. & Boyden, E. S. Optical imaging. Expansion microscopy. Science 347, 543–548 (2015). 10.1126/science.1260088

9 Tavakoli, M. R. et al. Light-microscopy-based connectomic reconstruction of mammalian brain tissue. Nature 642, 398–410 (2025). 10.1038/s41586-025-08985-1

10 Winding, M. et al. The connectome of an insect brain. Science 379, eadd9330 (2023). 10.1126/science.add9330

11 Zheng, Z. et al. A Complete Electron Microscopy Volume of the Brain of Adult Drosophila melanogaster. Cell 174, 730–743 e722 (2018). 10.1016/j.cell.2018.06.019

12 Dorkenwald, S. et al. Neuronal wiring diagram of an adult brain. Nature 634, 124–138 (2024). 10.1038/s41586-024-07558-y

13 Schlegel, P. et al. Whole-brain annotation and multi-connectome cell typing of Drosophila. Nature 634, 139–152 (2024). 10.1038/s41586-024-07686-5

14 Berg, S. et al. Sexual dimorphism in the complete Drosophila male central nervous system connectome. Cell 189, 5504–5526 e5515 (2026). 10.1016/j.cell.2026.08.015

15 Petkova, M. D. et al. A connectomic resource for neural cataloguing and circuit dissection of the larval zebrafish brain. bioRxiv (2025). 10.1101/2025.06.10.658982

16 Consortium, M. I. Functional connectomics spanning multiple areas of mouse visual cortex. Nature 640, 435–447 (2025). 10.1038/s41586-025-08790-w

17 Brand, A. H. & Perrimon, N. Targeted gene expression as a means of altering cell fates and generating dominant phenotypes. Development 118, 401–415 (1993).

18 Pfeiffer, B. D. et al. Tools for neuroanatomy and neurogenetics in Drosophila. Proc Natl Acad Sci U S A 105, 9715–9720 (2008). 10.1073/pnas.0803697105

19 Luan, H., Peabody, N. C., Vinson, C. R. & White, B. H. Refined spatial manipulation of neuronal function by combinatorial restriction of transgene expression. Neuron 52, 425–436 (2006). 10.1016/j.neuron.2006.08.028

20 Clements, J. et al. NeuronBridge: an intuitive web application for neuronal morphology search across large data sets. BMC Bioinformatics 25, 114 (2024). 10.1186/s12859-024-05732-7

21 Klapoetke, N. C. et al. Independent optical excitation of distinct neural populations. Nat Methods 11, 338–346 (2014). 10.1038/nmeth.2836

22 Mohammad, F. et al. Optogenetic inhibition of behavior with anion channelrhodopsins. Nat Methods 14, 271–274 (2017). 10.1038/nmeth.4148

23 Chen, T. W. et al. Ultrasensitive fluorescent proteins for imaging neuronal activity. Nature 499, 295–300 (2013). 10.1038/nature12354

24 Lin, A. et al. Network statistics of the whole-brain connectome of Drosophila. Nature 634, 153–165 (2024). 10.1038/s41586-024-07968-y

25 Masse, N. Y., Turner, G. C. & Jefferis, G. S. Olfactory information processing in Drosophila. Curr Biol 19, R700–713 (2009). 10.1016/j.cub.2009.06.026

26 Kurtovic, A., Widmer, A. & Dickson, B. J. A single class of olfactory neurons mediates behavioural responses to a Drosophila sex pheromone. Nature 446, 542–546 (2007). 10.1038/nature05672

27 Aso, Y. et al. The neuronal architecture of the mushroom body provides a logic for associative learning. Elife 3, e04577 (2014). 10.7554/eLife.04577

28 Schretter, C. E. et al. Cell types and neuronal circuitry underlying female aggression in Drosophila. Elife 9 (2020). 10.7554/eLife.58942

29 Wu, M. et al. Visual projection neurons in the Drosophila lobula link feature detection to distinct behavioral programs. Elife 5 (2016). 10.7554/eLife.21022

30 Ribeiro, I. M. A. et al. Visual Projection Neurons Mediating Directed Courtship in Drosophila. Cell 174, 607–621 e618 (2018). 10.1016/j.cell.2018.06.020

31 Hindmarsh Sten, T., Li, R., Otopalik, A. & Ruta, V. Sexual arousal gates visual processing during Drosophila courtship. Nature 595, 549–553 (2021). 10.1038/s41586-021-03714-w

32 Schretter, C. E. et al. Social state alters vision using three circuit mechanisms in Drosophila. Nature 637, 646–653 (2025). 10.1038/s41586-024-08255-6

33 Collie, M. F. et al. Specialized parallel pathways for adaptive control of visual object pursuit. Neuron 114, 1833–1847 e1839 (2026). 10.1016/j.neuron.2026.01.001

34 Taisz, I. et al. Generating parallel representations of position and identity in the olfactory system. Cell 186, 2556–2573 e2522 (2023). 10.1016/j.cell.2023.04.038

35 Asahina, K. Sex differences in Drosophila behavior: Qualitative and Quantitative Dimorphism. Curr Opin Physiol 6, 35–45 (2018). 10.1016/j.cophys.2018.04.004

36 Sato, K. & Yamamoto, D. Molecular and cellular origins of behavioral sex differences: a tiny little fly tells a lot. Front Mol Neurosci 16, 1284367 (2023). 10.3389/fnmol.2023.1284367

37 Zhou, C., Pan, Y., Robinett, C. C., Meissner, G. W. & Baker, B. S. Central brain neurons expressing doublesex regulate female receptivity in Drosophila. Neuron 83, 149–163 (2014). 10.1016/j.neuron.2014.05.038

38 Wang, K. et al. Neural circuit mechanisms of sexual receptivity in Drosophila females. Nature 589, 577–581 (2021). 10.1038/s41586-020-2972-7

39 Chiu, H. et al. A circuit logic for sexually shared and dimorphic aggressive behaviors in Drosophila. Cell 184, 507–520 e516 (2021). 10.1016/j.cell.2020.11.048

40 Deutsch, D. et al. The neural basis for a persistent internal state in Drosophila females. Elife 9 (2020). 10.7554/eLife.59502

41 Inagaki, H. K. et al. Optogenetic control of Drosophila using a red-shifted channelrhodopsin reveals experience-dependent influences on courtship. Nat Methods 11, 325–332 (2014). 10.1038/nmeth.2765

42 Hoopfer, E. D., Jung, Y., Inagaki, H. K., Rubin, G. M. & Anderson, D. J. P1 interneurons promote a persistent internal state that enhances inter-male aggression in Drosophila. Elife 4 (2015). 10.7554/eLife.11346

43 Koganezawa, M., Kimura, K. & Yamamoto, D. The Neural Circuitry that Functions as a Switch for Courtship versus Aggression in Drosophila Males. Curr Biol 26, 1395–1403 (2016). 10.1016/j.cub.2016.04.017

44 Rubin, G. M. et al. Networks of sexually dimorphic neurons that regulate social behaviors in Drosophila. Curr Biol 36, 4697–4716 e4697 (2026). 10.1016/j.cub.2026.08.013

45 Hindmarsh Sten, T., Li, R., Hollunder, F., Eleazer, S. & Ruta, V. Male-male interactions shape mate selection in Drosophila. Cell 188, 1486–1503 e1425 (2025). 10.1016/j.cell.2025.01.008

46 Asahina, K. et al. Tachykinin-expressing neurons control male-specific aggressive arousal in Drosophila. Cell 156, 221–235 (2014). 10.1016/j.cell.2013.11.045

47 Chiu, H. et al. Cell type-specific contributions to a persistent aggressive internal state in female Drosophila. Elife 12 (2025). 10.7554/eLife.88598

48 Lin, A. C., Bygrave, A. M., de Calignon, A., Lee, T. & Miesenbock, G. Sparse, decorrelated odor coding in the mushroom body enhances learned odor discrimination. Nat Neurosci 17, 559–568 (2014). 10.1038/nn.3660

49 Stimberg, M., Brette, R. & Goodman, D. F. Brian 2, an intuitive and efficient neural simulator. Elife 8 (2019). 10.7554/eLife.47314

50 Sapkal, N. et al. Neural circuit mechanisms underlying context-specific halting in Drosophila. Nature 634, 191–200 (2024). 10.1038/s41586-024-07854-7

